# Post-transcriptional regulation of glutamate metabolism of *Pichia pastoris* and development of a glutamate-inducible yeast expression system

**DOI:** 10.1101/2021.03.10.434748

**Authors:** Trishna Dey, Pundi N Rangarajan

## Abstract

*Pichia pastoris* harbours a unique glutamate utilization pathway in which glutamate dehydrogenase 2 (GDH2), aspartate aminotransferase 2 (AAT2) and phosphoenolpyruvate carboxykinase (PEPCK) catalyze the conversion of glutamate to α-ketoglutarate, oxaloacetate, and phosphoenolpyruvate respectively in the cytosol. GDH2 and PEPCK are glutamate-inducible enzymes and their synthesis is regulated post-transcriptionally by Rtg1p, a cytosolic basic helix-loop-helix protein via Rtg1p response elements located downstream of TATA box of *GDH2* and *PEPCK* promoters. Glutamate-inducible synthesis of PEPCK is abrogated in *Δgdh2* and *Δaat2*. α-ketoglutarate induces PEPCK synthesis in *Δgdh2* but not *Δaat2*. We propose that oxaloacetate derived from glutamate is the inducer of PEPCK synthesis. Enzymes of glutamate utilization pathway are synthesized during carbon starvation and they enable *P. pastoris* to overcome nutritional stress. Finally, green fluorescent protein can be synthesized efficiently from *GDH2* and *PEPCK* promoters using food-grade monosodium glutamate as inducer indicating that the post-transcriptional regulatory circuit described here can be exploited for the development of glutamate-inducible *P. pastoris* expression system.

## Introduction

Amino acids, in addition to their role as building blocks of proteins, play an important role in several metabolic reactions. In biosynthetic reactions, they directly participate as substrates or undergo deamination and contribute as ammonium ion and keto acids. Glutamate and glutamine serve as nitrogen donors for several nitrogenous compounds in the cell (Magasanik, 1993). In *Saccharomyces cerevisiae*, NADPH-dependent glutamate dehydrogenases (GDH1, GDH3) convert ammonia and α-ketoglutarate into glutamate. The latter combines with ammonia to generate glutamine in a reaction catalyzed by glutamine synthetase. A third, NAD-dependent GDH (GDH2) localized in the mitochondria degrades glutamate to α-ketoglutarate and ammonia and thus contributes to anaplerosis of Krebs cycle. Glutamate and glutamine contribute for 85% and 15% of total cellular nitrogen, respectively in *S. cerevisiae* (Miller and Magasanik, 1990) (Mara et al., 2018) (https://www.yeastgenome.org/locus/S000002374#protein) (Silao et al., 2020). In addition to being a nitrogen donor, Glutamate is also utilized as the sole source of carbon by *Pichia pastoris* (*a.k.a. Komagataella phafii*) and several other yeast species (Sahu and Rangarajan, 2016) (Freese et al., 2011). In *Scheffersomyces stipitis* and *P. pastoris*, GDH2 is required for the catabolism of glutamate and deletion of *GDH2* renders them defective in glutamate utilization (Sahu and Rangarajan, 2016) (Dey et al., 2018) (Freese et al., 2011). In addition to GDH2, aspartate amino transferases (AAT) encoded by *AAT1* and *AAT2*, malate dehydrogenases (MDH) encoded by *MDH1* and *MDH2* as well as the gluconeogenic enzyme, phosphoenolpyruvate carboxykinase (PEPCK) are also essential for glutamate utilization in *P. pastoris* (Sahu and Rangarajan, 2016) (Dey et al., 2018). Mxr1p, a zinc finger transcription factor and Rtg1p, a basic helix-loop-helix leucine zipper protein regulate the expression of key genes of glutamate utilization pathway of *P. pastoris* (Sahu and Rangarajan, 2016) (Dey et al., 2018). In *S. cerevisiae*, Rtg1p heterodimerizes with another basic helix-loop-helix leucine zipper protein known as Rtg3p and regulates the expression of specific nuclear genes in glutamate deficient cells with dysfunctional mitochondria, a phenomenon known as retrograde response (Jazwinski, 2014) (Butow and Avadhani, 2004). In *P. pastoris*, Rtg3p is absent and Rtg1p localizes to cytosol and functions as a post-transcriptional regulator of GDH2 and PEPCK, key enzymes of glutamate utilization pathway (Dey et al., 2018).

## Results and discussion

### Identification of Rtg1p response elements in *GDH2* and *PEPCK* promoters

We had identified several enzymes involved in the utilization of glutamate as the sole source of carbon in *P. pastoris* (Sahu and Rangarajan, 2016). Of these, GDH2 and PEPCK are glutamate-inducible enzymes and their synthesis from pre-existing mRNAs is regulated by the cytosolic, basic, helix-loop-helix leucine zipper protein, Rtg1p. (Dey et al., 2018). To further understand Rtg1p-mediated regulation of glutamate utilisation pathway of *P. pastoris*, genes encoding epitope tagged GDH2 and PEPCK were expressed from 1.0 kb of *GDH2* (*P_GDH2_*) and *PEPCK* (*P_PEPCK_*) promoters respectively (Supplementary Fig. 1) in *GS115* as well as mutant strains of *P. pastoris* (Supplementary Table 1). Cells were cultured in a medium containing yeast nitrogen base (YNB) and glucose (YNBD), glycerol (YNBG), acetate (YNBA), ethanol (YNBE) or glutamate (YNB Glu). GDH2 and PEPCK levels in cell lysates were examined by western blotting using anti-epitope tag antibodies while mRNA levels were quantified by qPCR. GDH2 was present only in cells cultured in YNB Glu while PEPCK was detected in cells cultured in YNBG, YNBE, YNBA and YNB Glu with highest levels in cells cultured in YNB Glu (Fig. 1A). Surprisingly, *GDH2* mRNA levels were comparable in cells utilizing glucose, glycerol, acetate, ethanol, and glutamate (Fig. 1B). *PEPCK* mRNA was undetectable during glucose metabolism and it was present in cells utilizing glycerol, ethanol, acetate, and glutamate (Fig. 1B). *PEPCK* mRNA levels were higher in cells utilizing glutamate and ethanol than other carbon sources (Fig. 1B). When *P_GDH2_* and *P_PEPCK_* were replaced by the promoter of the gene encoding glyceraldehyde-3-phosphate dehydrogenase (*P_GAPDH_*) (Supplementary Fig. 1), carbon source-specific synthesis of GDH2 and PEPCK was abrogated and they were expressed constitutively in cells cultured in different media (Fig. 1C). Thus, the absence of GDH2 and PEPCK when expressed from their own promoters under certain culture conditions is not due to their instability or degradation. To further understand carbon source-specific regulation of *GDH2* and *PEPCK*, *GFP* encoding green fluorescent protein was expressed from *P_GDH2_* and *P_PEPCK_*, GFP was visualized by live cell imaging while mRNA levels were examined by qPCR. GFP was expressed in a carbon source-specific manner (Fig. 1D) and the expression pattern was very similar to that of GDH2 and PEPCK. *GFP* mRNA levels were similar to *GDH2* and *PEPCK* mRNA levels (compare Fig. 1E with 1B). Analysis of GFP expression from *P_GDH2_* and *P_PEPCK_* in *GS115* and *Δrtg1* indicated that GFP protein levels were significantly downregulated in *Δrtg1* (Fig. 1G) without a corresponding decrease in their mRNA levels (Fig. 1H) indicating that Rtg1p regulates GFP synthesis from *P_GDH2_* and *P_PEPCK_* post-transcriptionally.

**Fig. 1.**
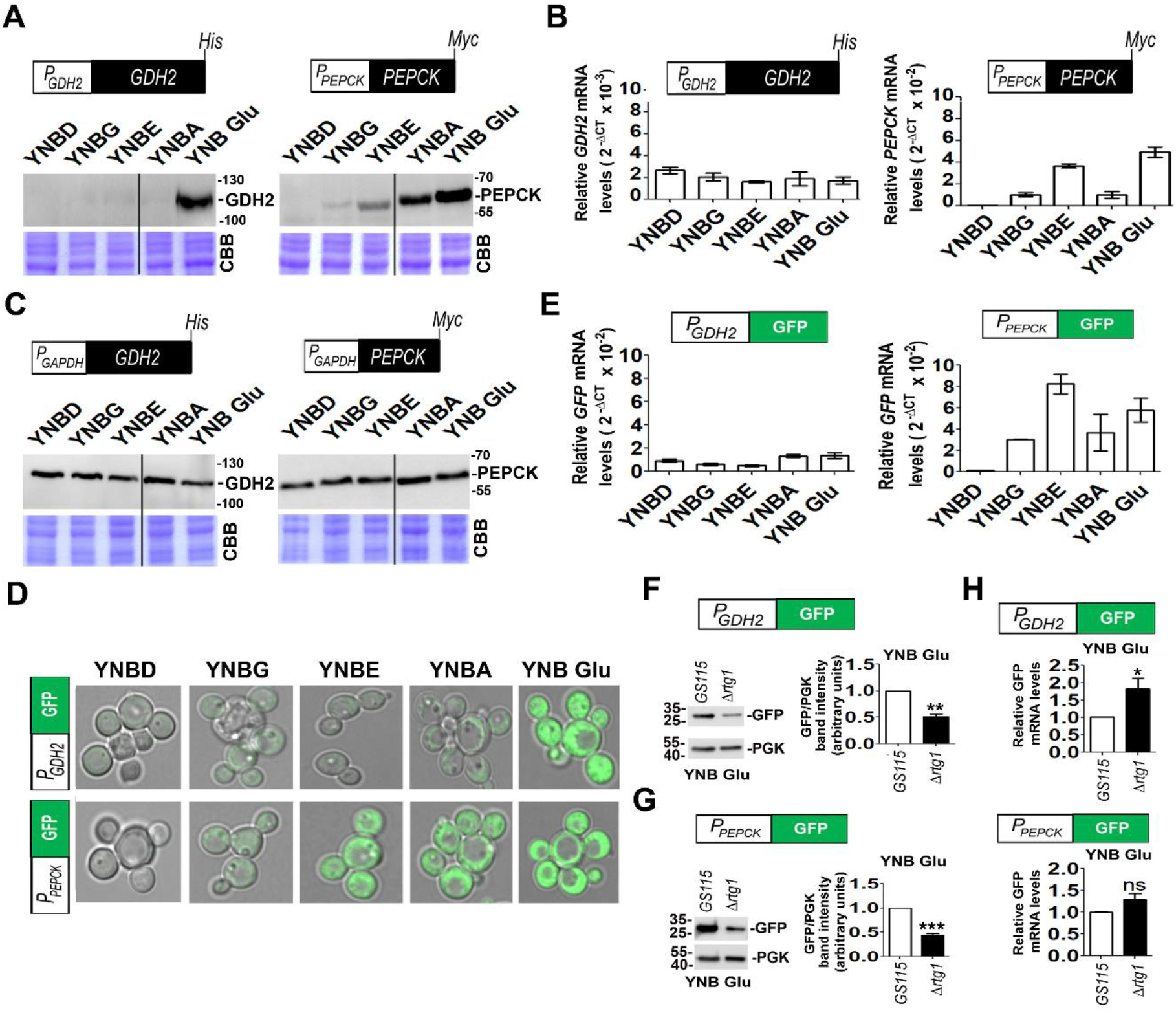
Carbon source specific, Rtg1p mediated post-transcriptional regulation of *GDH2* and *PEPCK*. **(A)** Epitope tagged GDH2 and PEPCK were expressed from their own promoters (schematically represented) and protein levels were examined by western blotting using anti epitope tag antibodies in the whole cell lysates of cells utilizing different carbon sources. YNBD, Yeast nitrogen base (YNB) dextrose; YNBG, YNB glycerol; YNBE, YNB ethanol; YNBA, YNB acetate and YNB Glu, YNB glutamate. **(B)** Expression profile of *GDH2* and *PEPCK* mRNAs in cells utilizing different carbon compounds by qPCR. **(C)** Western blot analysis of epitope tagged GDH2 and PEPCK proteins synthesized from a heterologous promoter encoding *GAPDH* (*P_GAPDH_*). **(D)** GFP expression profile from *GDH2* (*P_GDH2_*) and *PEPCK* (*P_PEPCK_*) promoters in cells utilizing different carbon sources by live cell confocal imaging. **(E)** qPCR analysis of *GFP* mRNA expressed from *P_GDH2_* and *P_PEPCK_* in cells cultured in media containing different carbon sources. **(F, G)** Western blot analysis of GFP synthesized from *P_GDH2_* and *P_PEPCK_* in *GS115* (WT) and *Δrtg1* cells cultured in YNB Glu. The intensities of the protein bands were quantified, normalised to control (PGK) and plotted relative to *GS115*. PGK, phoshoglycerate kinase served as loading control. **(H)** qPCR analysis of GFP mRNA levels expressed from *P_GDH2_* and *P_PEPCK_* in *Δrtg1* relative to *GS115* cultured in YNB Glu medium. Error bar denotes mean±S.D. (Biological replicates, n = 3). P value is obtained from Student’s t-test and is mentioned on the bar of each figure: * *P* <0.05; ** *P* <0.005; *** *P* <0.0005, ns not significant. CBB is the Coomassie Brillaint Blue R-stained SDS-polyacrylamide gel **(A, C)** serves as a loading control. Numbers **(A,C,F and G)** indicate protein molecular weight markers (kDa).

The results thus far indicate that Rtg1p is a post-transcriptional regulator of *GDH2* and *PEPCK* suggesting that upstream promoter sequences of *P_GDH2_* and *P_PEPCK_* which act as binding sites for transcription factors are unlikely to have a role in Rtg1p-mediated synthesis of GDH2 and PEPCK. We, therefore, focused our attention on promoter sequences downstream of TATA box upto the initiation codon (ATG). In yeast promoters, transcription start site (TSS) is located 40-120 bp downstream of TATA box and it corresponds to the first base of mRNA (McMillan et al., 2019). The region between the TSS and the ATG in the promoter corresponds to the 5’ untranslated region (5’ UTR) of mRNA which often contains cis-acting elements referred to as riboswitches that bind to small molecules and regulate translation of downstream open reading frames (Garst et al., 2011). Since TSSs of *P. pastoris GDH2, PEPCK* and *GAPDH* are not characterized, we examined the role of downstream promoter region (DPR) located between the putative TATA box and the initiation codon of *P_GDH2_* and *P_PEPCK_* in the Rtg1p-dependent, post-transcriptional regulation (Fig. 2a). DPR of *P_GAPDH_* (−99 to −1 bp) was replaced with that of *P_GDH2_* (−86 to −1 bp) or *P_PEPCK_* (−58 to −1 bp) (Fig. 2A) and synthesis of GDH2 and PEPCK from these chimeric promoters was examined in lysates of *GS115* and *Δrtg1* cultured in YNB Glu medium by western blotting with anti-epitope tag antibodies. As expected, GDH2 and PEPCK levels were comparable in *GS115* and *Δrtg1* when *P_GAPDH_* was used (Fig. 2B, C). However, when DPR of *P_GAPDH_* was replaced with that of *P_GDH2_* or *P_PEPCK_*, GDH2 and PEPCK synthesis became Rtg1p-dependent resulting in the down regulation of protein but not mRNA levels in *Δrtg1* (Fig. 2D, E). Thus, the DPRs harbour the cis-acting elements required for Rtg1p-mediated synthesis of GDH2 and PEPCK.

**Fig. 2.**
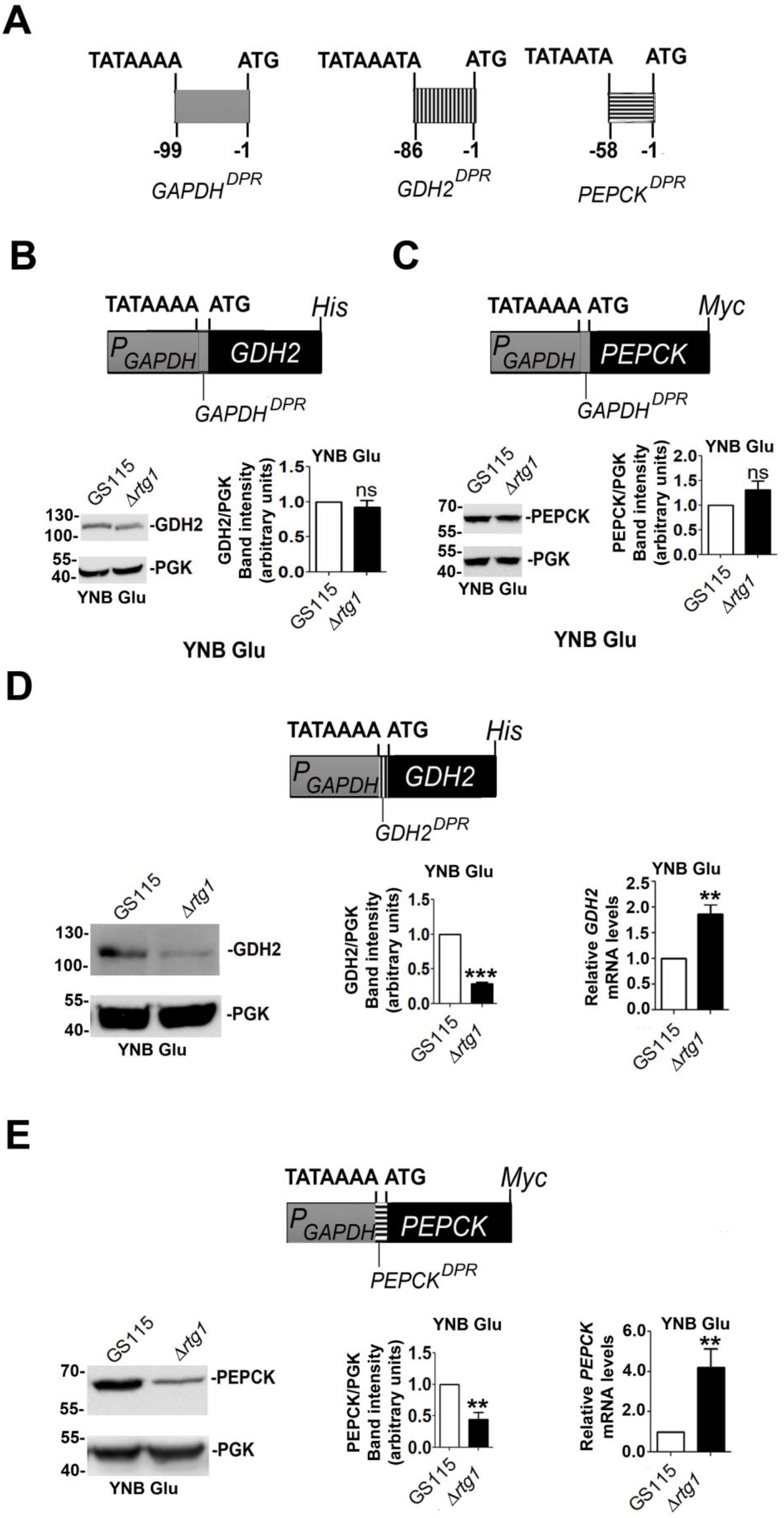
Identification of Rtg1p response element within the downstream promoter region (DPR) of *P_GDH2_* and *P_PEPCK_*. **(A)** Schematic representation of DPR between the TATA box and the initiation codon of *P_GAPDH_*, *P_GDH2_* and *P_PEPCK_*. **(B)** Western blot analysis of His tagged GDH2 and **(C)** Myc tagged PEPCK synthesied from *P_GAPDH_* containing *GAPDH^DPR^* in *GS115* and *Δrtg1* cells cultured in YNB Glu medium. **(D)** Analysis of His tagged GDH2 and **(E)** Myc tagged PEPCK expression from chimeric *P_GAPDH_* containing *GDH2^DPR^* and *PEPCK^DPR^*, respectively, in *GS115* and *Δrtg1* cells cultured in YNB Glu medium. Protein and mRNA levels were analysed by western blotting and qPCR respectively. The intensities of the protein bands were quantified, normalised to control (PGK) and plotted relative to *GS115.* Error bar denotes mean±S.D. (Biological replicates, n = 3). P value is obtained from Student’s t-test and is mentioned on the bar of each figure: * *P* <0.05; ** *P* <0.005; *** *P* <0.0005, ns not significant. Numbers in **(B-E)** indicate molecular weight marker (kDa).

### Conversion of glutamate to oxaloacetate is essential for glutamate-inducible synthesis of PEPCK

GDH2 and PEPCK are abundant proteins and can be readily visualized in Coomassie Brilliant Blue-stained SDS polyacrylamide gels of lysates of *GS115* cultured in YNB Glu (Fig 3A) (Dey et al., 2018). These protein bands were identified as GDH2 and PEPCK by Mass spectrometry in an earlier study (Dey et al., 2018). As expected, GDH2 and PEPCK were down regulated in *Δrtg1* cultured in YNB Glu. (Fig. 3A). However, GDH2 was down regulated in *Δpepck* and PEPCK was downregulated in *Δgdh2* (Fig. 3A). These results were confirmed by examining the levels of epitope tagged GDH2 and PEPCK by western blotting of cell lysates using anti-epitope tag antibodies (Fig. 3B). *GDH2* and *PEPCK* mRNAs are not downregulated (Fig. 3C) and downregulation of PEPCK in *Δgdh2* during glutamate metabolism was observed only during the utilization of glutamate but not acetate and ethanol (Fig. 3D). The first three reactions of glutamate catabolism catalyzed by GDH2, AAT2 and PEPCK result in the generation of α-ketoglutarate (Akg), oxaloacetate (Oaa) and phosphoenolpyruvate in the cytosol (Fig. 3E) (Sahu and Rangarajan, 2016). We hypothesized that glutamate may not be the direct inducer of PEPCK synthesis. To test this hypothesis, we examined the ability of Akg to induce GDH2 and PEPCK synthesis. Akg readily induced the synthesis of PEPCK but not GDH2 (Fig. 3F). Akg also induced GFP expression from *P_PEPCK_* but not *P_GDH2_* (Fig. 3G) suggesting that inducers of GDH2 and PEPCK synthesis are different. Since the synthesis of Akg is impaired in *Δgdh2*, we examined Akg-inducible PEPCK synthesis in Δgdh2. *GS115* and *Δgdh2* expressing Myc-tagged PEPCK were cultured in YNB Glu or YNB Akg and PEPCK protein levels were examined by western blotting. Only Akg but not glutamate induced PEPCK synthesis in *Δgdh2* (Fig. 3H). Since Akg is converted to Oaa by AAT2 and Oaa is the substrate of PEPCK, we examined the ability of Oaa to induce PEPCK synthesis. However, PEPCK synthesis was not induced in cells cultured in YNB Oaa (data not shown). We therefore examined the ability of Oaa synthesized intracellularly via AAT2-catalyzed reaction to induce PEPCK synthesis. Both glutamate- and Akg-inducible synthesis of PEPCK was abrogated in *Δaat2* (Fig. 3I) suggesting that Oaa is the inducer of PEPCK synthesis. Based on these results, we propose a model for the post-transcriptional regulation of *P. pastoris* glutamate utilization pathway wherein *GDH2^DPR^* and *PEPCK^DPR^* contain cis-acting elements named Rtg1p response elements (RRE) between the transcription start site and initiation codon. Transcription results in the generation of *GDH2* and *PEPCK* mRNAs harbouring the RREs in their 5’ UTRs (5’ UTR^RRE^). Translation of *GDH2* mRNA is facilitated by the interaction of glutamate and Rtg1p with the 5’ UTR^RRE^ of *GDH2* mRNA while PEPCK synthesis requires interactions amongst Rtg1p and oxaloacetate and 5’ UTR^RRE^of *PEPCK* mRNA (Fig. 3J).

**Fig. 3.**
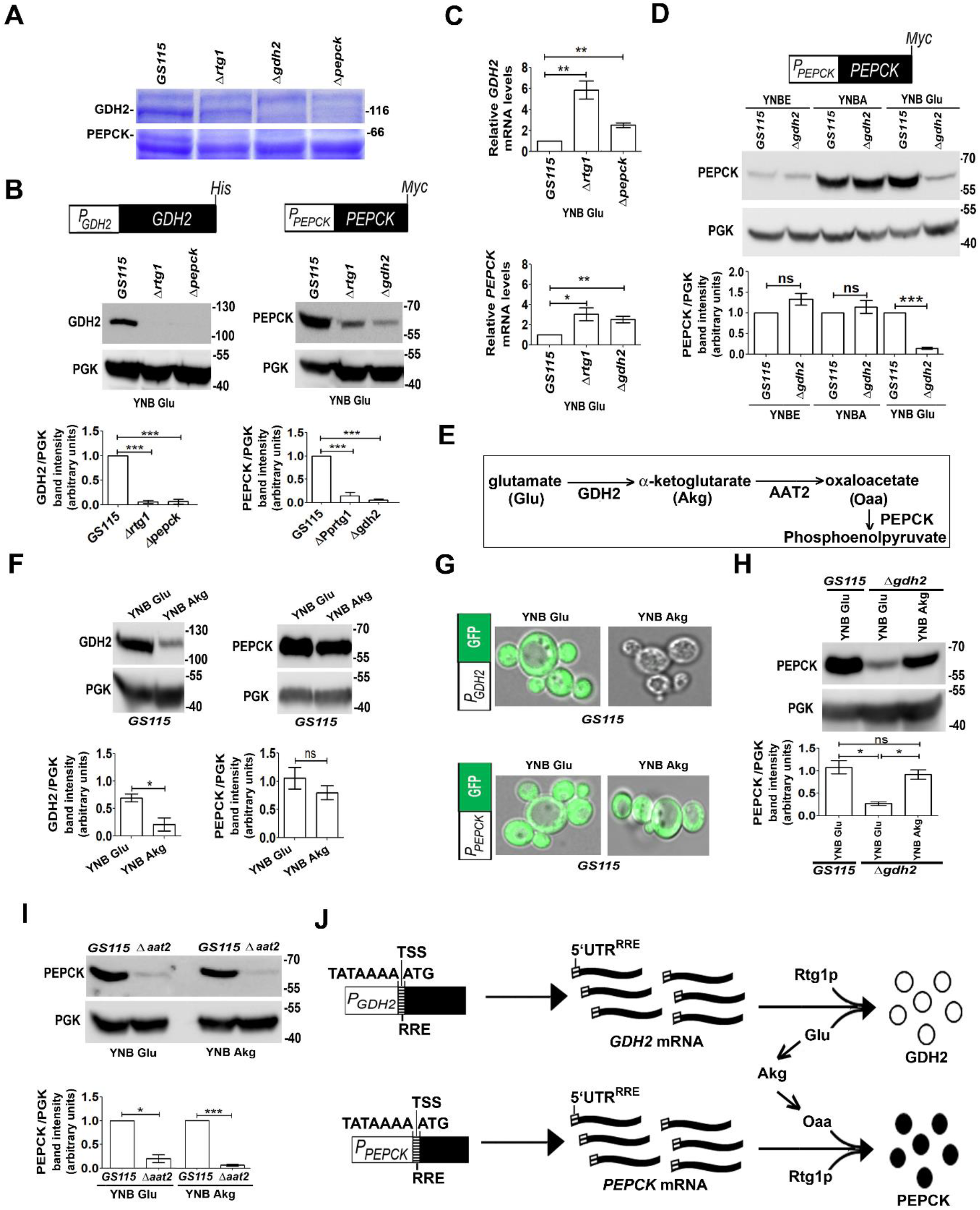
Identification of oxaloacetate as the inducer of PEPCK synthesis during glutamate metabolism. **(A)** Protein profile of lysates of *GS115, Δrtg1, Δgdh2 and Δpepck* cultured in YNB Glu medium. Proteins were resolved on SDS-polyacrylamide gel and stained with Coomassie Brilliant Blue R. Differential expression of GDH2 and PEPCK in *GS115, Δrtg1, Δgdh2 and Δpepck* is shown. Protein bands in *GS115* lane were identified as GDH2 and PEPCK by mass spectrometry in an earlier study (Dey et a., 2018). **(B)** Schematic representation of epitope tagged constructs of *GDH2* and *PEPCK* expressing in *GS115, Δrtg1, Δgdh2 and Δpepck* from 1 kb of *P_GDH2_* and *P_PEPCK_*, respectively. Western blot analysis of GDH2 and PEPCK levels in the whole cell protein lysates of cells cultured in YNB Glu medium using anti-His and anti-Myc antibodies, respectively. PGK served as loading control. The intensities of the protein bands were quantified, normalised to control (PGK) and plotted relative to *GS115*. (**C)** qPCR analysis of *GDH2* and *PEPCK* mRNA levels in *Δpepck and Δgdh2,* respectively, relative to *GS115.* **(D)** Analysis of Myc tagged PEPCK levels in the whole cell protein lysates of *GS115* and *Δgdh2* cultured in YNB Glu, YNBE and YNBA by western blotting using anti-Myc antibody. PGK served as loading control. The intensities of the protein bands were quantified, normalised to control (PGK) and plotted relative to *GS115*. **(E)** Schematic representation of first three reactions of glutamate utilisation pathway of *P. pastoris* catalyzed by the cytosolic enzymes GDH2, AAT2 and PEPCK. **(F)** Analysis of His tagged GDH2 and Myc tagged PEPCK expression in *GS115* cultured in YNB Glu and YNB Akg by western blotting. The intensities of the protein bands were quantified, normalised to control (PGK) and plotted relative to *GS115* cultured in YNB Glu. **(G)** Analysis of GFP expression from *P_GDH2_* and *P_PEPCK_* in *GS115* cultured in YNB Glu and YNB Akg by live cell confocal imaging**. (H)** Western blot analysis of Myc tagged PEPCK levels in *Δgdh2* cultured in YNB Akg medium compared to the levels in *GS115* and *Δgdh2* cultured in YNB Glu using anti-Myc antibody. PGK served as loading control. The intensities of the protein bands were quantified, normalised to control (PGK) and plotted relative to *GS115* cultured in YNB Glu. **(I)** Analysis of expression of Myc tagged PEPCK in *GS115* and *Δaat2* cultured in YNB Glu and YNB Akg by western blotting using anti-Myc antibody. PGK served as loading control. The intensities of the protein bands were quantified, normalised to control (PGK) and plotted relative to *GS115*. **(J)** A model for the post-transcriptional regulation of glutamate utilisation pathway of *P. pastoris*. The downstream promoter region (DPR) of *GDH2* and *PEPCK* promoters harbour putative Rtg1p response elements (RREs) between the transcription start site (TSS) and initiation codon. Upon transcription, RREs become part of the 5’ UTRs of mRNAs. Translation of *GDH2* mRNA requires complex interactions involving 5’ UTR^RRE^, glutamate (Glu) and Rtg1p. Translation of *PEPCK* mRNA requires complex interactions involving 5’ UTR^RRE^, oxaloacetate (Oaa) and Rtg1p. Akg is α-ketoglutarate. Error bar denotes mean±S.D. (Biological replicates, n = 3). *P* value is obtained from Student’s t-test and is mentioned on the bar of each figure: * *P* <0.05; ** *P* <0.005; *** *P* <0.0005, ns not significant. Numbers in (**A, B** and **D)** and (**F, H and I)** indicate molecular weight marker (kDa).

### Glutamate catabolism confers tolerance to carbon starvation

The results presented thus far demonstrate that amino- or keto-acids added to the culture medium can induce the translation of *GDH2* and *PEPCK* mRNAs in *P. pastoris*. GDH2 is known to funnel the carbon skeletons of glutamate into the TCA cycle for energy production under carbon limiting conditions (Miyashita and Good, 2008). Yeasts such as *P. pastoris* are likely to experience nutrient starvation in their natural habitats and they evolve several strategies to survive under these conditions. We reasoned that intracellular metabolites such as amino- and keto-acids generated during carbon starvation may induce the synthesis of GDH2 and PEPCK to facilitate energy generation and gluconeogenesis. *P. pastoris* cells expressing epitope-tagged GDH2 and PEPCK were cultured in nutrient-rich medium and then shifted to starvation medium which is completely devoid of an exogenous carbon source. Synthesis of GDH2 and PEPCK proteins from pre-existing mRNAs was induced within 6 h of carbon starvation (Fig. 4A, B). Carbon starvation results in the activation of autophagy which in turn results in the generation metabolites needed for energy generation, biosynthetic reactions, homeostasis and survival (Iwama and Ohsumi1, 2019) (Adachi et al., 2017) (Weber et al., 2020). To examine whether the translation of *GDH2* and *PEPCK* mRNAs is triggered by the metabolites generated upon induction of autophagy during carbon starvation, we treated cells with methionine, a known inhibitor of non-nitrogen starvation induced autophagy (Sutter et al., 2013) and analyzed GDH2 and PEPCK protein levels by western blotting. Synthesis of PEPCK but not GDH2 was impaired by methionine treatment (Fig. 4C). Thus, inducers of GDH2 and PEPCK synthesis are generated by autophagy-independent and -dependent pathways during carbon starvation, respectively. We next compared the viability of *GS115, Δgdh2 and Δpepck* during carbon starvation by cell survival assay. *GS115* expressing GDH2 and PEPCK exhibited greater viability than *Δgdh2* and *Δpepck* when observed over a period of two weeks of carbon starvation (Fig. 4D, E). The decrease in viability was much more pronounced in case of *Δpepck* than *Δgdh2.* Thus, starvation-induced synthesis of GDH2 and PEPCK from pre-existing mRNAs enables *P. pastoris* to rapidly respond to extreme carbon starvation conditions and overcome nutritional stress.

**Fig. 4.**
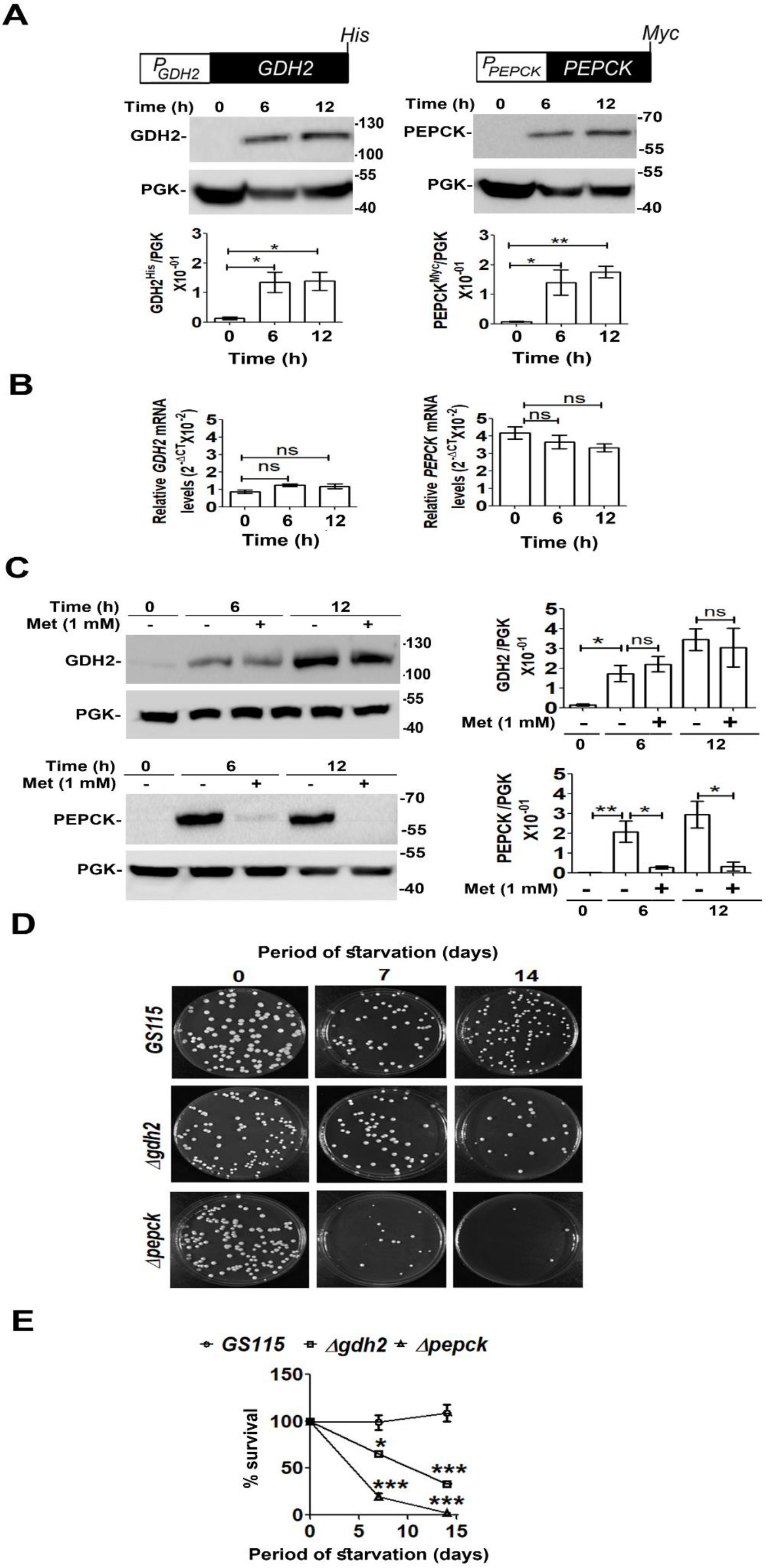
GDH2 and PEPCK confer survival advantage to *P. pastoris* during carbon starvation. **(A)** Analysis of expression of epitope tagged GDH2 and PEPCK in *GS115* subjected to carbon starvation for 6-12 h. GDH2 and PEPCK levels were examined by western blotting of cell lysates using anti-His and anti-Myc antibodies, respectively. PGK served as loading control. The intensities of the protein bands were quantified, normalised to control (PGK) and plotted relative to 0 h of starvation. **(B)** qPCR analysis of *GDH2* and *PEPCK* mRNA levels in *GS115* cultured in starvation medium. **(C)** Western blot analysis of GDH2 and PEPCK protein levels in cells cultured in starvation medium in the presence or absence of 1 mM methionine. PGK served as loading control. The intensities of the protein bands were quantified, normalised to control (PGK) and plotted relative to 0 h of starvation. **(D)** Analysis of viability *GS115, Δgdh2* and *Δpepck* upon carbon starvation by cell survival assay. At indicated time points, nutrient-deprived cultures were removed and plated on YPD agar containing yeast extract (1%), peptone (2%) and dextrose (1%). **(E)** Quantitation of cell survival (**D**). Number of colonies on the agar plates were counted and percent colony forming unit was plotted as percent survival. Error bar denotes mean±S.D. (Biological replicates, n = 3). *P* value is obtained from Student’s t-test and is mentioned on the bar of each figure: * *P* <0.05; ** *P* <0.005; *** *P* <0.0005, ns not significant. Numbers in **A** and **C** indicate molecular weight marker (kDa).

### A glutamate-inducible *P. pastoris* expression system

The promoter of *AOXI* (*P_AOXI_*) encoding alcohol oxidase I is widely used for methanol-inducible production of heterologous proteins in *P. pastoris* (Hartner and Glieder, 2006) (Cereghino and Cregg, 2000). Methanol is toxic and flammable compound, needs special handling and is not recommended to produce certain edible and medical products. Monitoring methanol concentration in *P. pastoris* cultures is necessary to prevent accumulation of formaldehyde and H_2_O_2_, by-products of methanol metabolism, to toxic levels necessitating the development of methanol-free *P_AOXI_*-based expression systems (Vogl et al., 2018) (Wang et al., 2017) (Shen et al., 2016) (Prielhofer et al., 2013). Thus, there is a need to develop expression systems which avoid methanol but maintain the high productivity of *P. pastoris*. Food grade monosodium glutamate (MSG) is more readily soluble in water than glutamate, generally regarded as safe (GRAS) by USFDA, available in large quantities, and used as a flavour enhancer in several Asian cuisines (https://www.amazon.in/Ajinomoto-Monosodium-Glutamate-Umami-Seasoning/dp/B00IH28XDE) (https://www.fda.gov/food/food-additives-petitions/questions-and-answers-monosodium-glutamate-msg). MSG readily induced GFP expression from *P_GDH2_* and *P_PEPCK_* (Fig. 5A). *P. pastoris* strain expressing GFP from *P_AOXI_* was constructed and GFP synthesis from methanol-inducible *P_AOXI_* was compared with that from MSG-inducible *P_GDH2_* and *P_PEPCK_*. Cells were cultured overnight in shake flasks in YNBD medium and then shifted to YNB and methanol (YNBM) or YNB Glu. After 24 h, cells were lysed and GFP levels were examined by western blotting using anti-GFP antibody (Fig. 5B). In another experiment, whole cell lysates were incubated with GST-tagged anti-GFP nanobodies (Katoh et al., 2015), proteins were pulled down using glutathione resin, and visualized on SDS polyacrylamide gel by Coomassie Brilliant Blue staining (Fig. 5C). Western blot and Coomassie blue-stained SDS gel were scanned and GFP was quantitated. The results indicate that GFP expression from *P_GDH2_* and *P_PEPCK_* is 65% and 90% respectively of that from P_*AOXI*_ (Fig. 5D). GFP was also quantified using a standard curve prepared using known quantities of BSA. GFP expression from *P_GDH2_* and *P_PEPCK_* and P_*AOXI*_ was 8.3, 12.6 and 14.4 μg per 1.2 ml of culture (Fig. 5E).

**Fig. 5.**
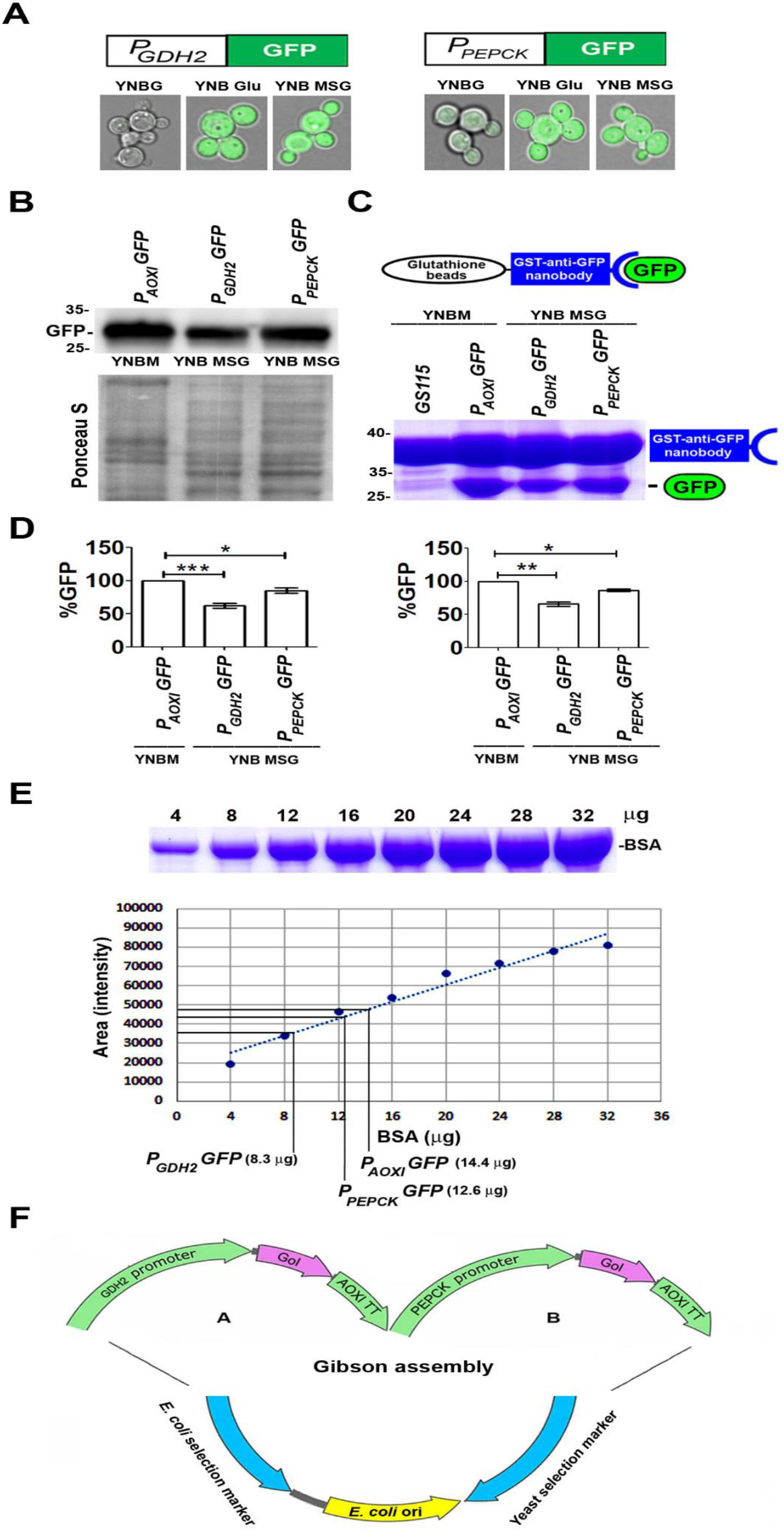
A monosodium glutamate (MSG)-inducible *Pichia pastoris* expression system. **(A)** GFP expression profile from *P_GDH2_* and *P_PEPCK_* in cells cultured in media containing either glycerol, glutamate or MSG as the sole source of carbon by live cell confocal imaging. **(B)** Comparative analysis of GFP expression from *P_AOX1_*, *P_GDH2_* and *P_PEPCK_* in cells cultured in YNBM (*P_AOX1_*) and YNB MSG (*P_GDH2_* and *P_PEPCK_*) by western blotting. **(C)** GST tagged anti-GFP nanobody mediated pull down assay. GST-tagged anti-GFP nanobodies obtained from *E. coli* lysates (Katoh et al., 2016) were incubated with glutathione-sepharose beads and *P. pastoris* cell lysates as indicated. After washing, proteins bound to glutathione-sepharose beads were resolved on SDS-PAGE and visualized by Coomassie Brilliant Blue staining. **(D)** Quantitation of GFP levels in **(B)**(left panel) and (**C)**(right panel). The intensities of the protein bands were quantified and normalised to GFP expressed from *P_AOX1_* and YNBM cultured cells. Error bar denotes mean±S.D. (Biological replicates, n = 3). *P* value is obtained from Student’s t-test and is mentioned on the bar of each figure: * *P* <0.05; ** *P* <0.005; *** *P* <0.0005, ns not significant. **(E)** Estimation of amount of GFP in (**C)** using BSA standard curve generated from known concentrations of BSA. **(F)** Pictorial representation of a glutamate inducible expression vector containing two glutamate inducible expression cassettes. Gene of interest (GoI) can be expressed from *GDH2* and *PEPCK* promoters simultaneously in the same *P. pastoris* strain maximising recombinant protein yield. Numbers in **(B)** and **(C)** indicate protein molecular weight markers (kDa). Ponceau S-stained blot **(B)** serves as loading control.

The efficiency of the glutamate-inducible expression system described here can be further improved by expressing the gene of interest from both *P_GDH2_* and *P_PEPCK_* (Fig. 5F). The combined strength of *P_GDH2_* and *P_PEPCK_* is likely to be more than that of *P_AOXI_*. While GDH2 is expressed only in cells utilizing glutamate, PEPCK is synthesised in cells utilizing glutamate as well as ethanol, acetate and glycerol albeit at varying levels. (Fig. 1A-D). Thus, by choosing an appropriate carbon source, one can express a heterologous gene at low, moderate, or high levels from *PEPCK* promoter which can be of great importance in expressing proteins that are cytotoxic at high levels as well as expressing proteins of a metabolic pathway at variable levels during metabolic engineering. The high-density fermentation strategy currently being employed for methanol-inducible production of recombinant proteins can be readily adapted for glutamate-inducible expression with only minor modifications. The expression system described here is unique as recombinant protein production involves post-transcriptional regulation. *GDH2*- and *PEPCK* promoter-based *P. pastoris* expression system will be useful for both academic research and industrial applications.

Taken together, this study demonstrates that the glutamate utilization pathway is of great physiological significance. Glutamate-inducible synthesis of GDH2 and PEPCK is regulated by Rtg1p in conjunction with their respective substrates and RREs. The fact that the translation of *GDH2* and *PEPCK* is highly regulated and they express at high levels in cells cultured in YNB Glu medium has been exploited for the development of a novel, glutamate-inducible *P. pastoris* expression system for the synthesis of heterologous proteins as an alternative to the methanol-inducible expression system.

## Materials and methods

### Media and culture conditions

A single colony of yeast cells was inoculated from agar (2.0%) plates containing YPD (1.0% yeast extract, 2.0% peptone, 2.0% glucose) into YPD liquid medium and grown overnight at 30°C in an orbital shaker at 180 rpm. Cells were washed with sterile distilled water, atleast twice, and shifted to different minimal media containing 0.17% yeast nitrogen base (YNB) without amino acids and with 0.5% ammonium sulphate supplemented with 2.0% glucose (YNBD), 2.0% glycerol (YNBG), 1.0% ethanol (YNBE), 2.0% acetate (YNBA), 1.0% methanol (YNBM) or 1.0% glutamate (YNB Glu).

### Yeast and bacterial strains

Yeast strains used in this study are listed in Supplementary Table1. Yeast cells were transformed by electroporation (Gene Pulser, Bio-Rad, CA). *Escherichia coli DH5α* strain was used for cloning of recombinant plasmids and transformation was done by CaCl_2_ method.

### Antibodies and other reagents

Oligonucleotides were purchased form Sigma-Aldrich (Bangalore, India). Mouse anti-Myc tag antibody was purchased from Merck Millipore (Bangalore, India). Mouse anti-GFP was purchased from Santa Cruz Biotechnology Inc. (Santa Cruz, CA) and mouse anti-His tag antibodies were purchased from cell signalling and technology (Denvers, MA). Anti-phosphoglyerate kinase (PGK) antibody was generated by immunizing rabbit with purified *P. pastoris* PGK protein. Restriction enzymes and T4 DNA ligase were purchased from New England Biolabs (Ipswich, MA). DNA polymerases were purchased from GeNei (Bangalore, India) and Thermo Fisher Scientific (Waltham, MA).

### Total RNA isolation, Reverse Transcription-PCR (RT-PCR) and Quantitative PCR (qPCR)

Total RNA was isolated from *P. pastoris* by hot-phenol method (Schmitt et al., 1990). Yeast cells grown in a specified medium were harvested by centrifugation (3000 × g for 5 min) at room temperature. Cells were resuspended in 400 μl of AE buffer (50 mM sodium acetate, pH 5.3 and 10 mM EDTA, pH 8). To this 40 μl of 10% SDS was added and vortexed briefly. After vortexing, 500 μl of AE buffer-saturated phenol was added, vortexed and heated at 65°C for 4 min. The tubes were then snap chilled in liquid nitrogen for 1 min and centrifuged (10000 × g for 15 min) for the separation of aqueous and organic phases. The aqueous phase was carefully transferred to a fresh tube followed by phenol-chloroform extraction in which equal volume of phenol-chloroform (1:1) was added, vortexed briefly and centrifuged (10000 × g for 5 min) at room temperature. To the upper aqueous phase, 40 μl of 3 M sodium acetate (pH 5.3) and 2.5 volumes of chilled 100% ethanol was added, mixed thoroughly by inverting and incubated at −20°C for 30 min for the precipitation of RNA. RNA pellet was obtained by centrifugation at 10000 × g for 15 min at 4°C. RNA pellet was washed with chilled 80% ethanol, air dried and resuspended in water. For quantitation, absorbance of RNA was recorded at 260 nm. cDNA was prepared using 1 μg of DNase treated RNA followed by q-PCR using StepOnePlus Real-Time PCR system (Thermo Fisher Scientific).

### Western blotting

Whole cell protein extracts were made by either glass bead lysis or trichloro- acetic acid (TCA) precipitation method. For TCA precipitation, briefly 5.0 OD_600_ units of *P. pastoris* cells cultured in specified medium were centrifuged (Hettich, 3000 × g for 5 min) at room temperature, resuspended in 1 ml 20% TCA, vortexed thoroughly and centrifuged again (3000 × g for 5 min) at room temperature. TCA was removed completely and to the pellet 200 μl 20% TCA was added. Acid washed 0.5 mm glass beads (Biospec Products) were added in 1:1 ratio (w/w) and vortexed (at highest setting) 4 times with intermittent chilling on ice. The slurry was aspirated in fresh microfuge tube and TCA was removed by centrifugation (10000 × g at 4°C for 15 min). Brief spin was given again to remove left over TCA. 1X Laemmli buffer (50 mM Tris-HCl, pH 6.8, 2% SDS, 0.1% bromophenol blue, 10% glycerol and 100 mM β-mercarptoethanol) was added to the pellet, vortexed, heated for 5 min at 95°C and vortexed again. Final whole cell protein lysate was extracted from left over debris by centrifugation (10000 × g at 4°C for 10 min). Lysate volume corresponding to 0.5 OD_600_ units of cells was resolved on SDS-PAGE. For Glass bead mediated lysis, yeast cells were resuspended in lysis buffer (1:1 w/v) containing 20 mM Tris (pH 8.0), 400 mM NaCl, 10 mM MgCl2, 10 mM EDTA (pH 8.0), 10% glycerol, 7 mM β-mercaptoethanol and protease inhibitor cocktail (Roche complete, EDTAfree Protease Inhibitory Cocktail tablets, Manheim, Germany). Acid washed 0.5 mm glass beads (Biospec Products) were added in 1:1 ratio (w/w) and vortexed 10 times using a vortex mixer (at highest setting) with intermittent chilling on ice. Cell debris were removed by centrifugation (10000 × g at 4°C). Proteins were estimated using Bradford reagent (BioRad) and resolved on SDS-PAGE, electroblotted onto a 0.22 μm PVDF membrane using transfer buffer (39 mM glycine, 48 mM Tris (pH 8.0), 20% methanol). The membrane was blocked in 5% non-fat milk for 1 h, prepared in TTBS (25 mM Tris (pH 8.0), 0.1% Tween 20 and 125 mM NaCl) followed by three washes in TTBS for 5 min. Blots were sequentially incubated in TTBS containing antibodies of appropriate dilution, raised against a specific protein or anti-epitope tag antibodies, for 1h either at room temperature or overnight at 4°C and washed in TTBS (thrice for 5 min). Primary antibodies were detected by HRP-conjugated anti-rabbit/anti-mouse IgG (1:10000 dilution, 1 h and 4°C incubation). Proteins were detected using Immobilon Western Chemiluminescent HRP substrate (Millipore MA) as per the manufacturer’s instructions.

### Live cell imaging

For live cell imaging, 3 μl of growing culture was layered on top of an agarose pad and allowed to settle for 1 min. Finally, on top of the cell suspension coverslip was placed and sealed with transparent nail paint. GFP expression in cells was then visualised using confocal microscope (Olympus FLUOVIEW FV3000). To make agarose pad, glass slides were wiped thoroughly and both ends were fixed with scotch tape. 30 μl of molten 1.0% agarose (in 1X PBS) was placed on top of the glass slide into a round droplet. Another clean slide was placed on top of the agarose droplet and pressed gently to make the flattened agarose pad. This entire set up of the agarose pad was lifted carefully and placed on top of a precooled metal block for 1 min. After the solidification of agarose, the glass slides were separated by gentle sliding motion leaving the agarose pad stuck to one of the two slides which is used further.

### Cell viability assay

From the frozen stock, strains were streaked onto YPD plates and incubated at 30°C for 2-3 days. Single colony was picked and inoculated in YPD medium. Cells were grown in YPD at 180 rpm, 30°C till late stationary phase (2 days). YPD grown late stationary phase cells were transferred to starvation medium i.e., 0.17% YNB without amino acids and with 0.5% ammonium sulphate supplemented with 20 μg/ml of histidine with initial OD_600_ of ~0.5 per ml of media. At specified time points, OD_600_ was recorded and adjusted to ~0.1 per ml. 1:10 serial dilutions were made and 50 μl of 10^−3^ dilution was spread on YPD plates. Plates were incubated at 30°C for 2-3 days and colony forming unit per ml (CFU/ml) was calculated as a readout for cell survival. CFU per ml at day zero was considered 100%.

### Generation of *P. pastoris GS115* and *Δrtg1* expressing *GDH2^His^* and *PEPCK^Myc^* from *P_GDH2_* and *P_PEPCK_*, respectively, *Δgdh2* expressing *PEPCK^Myc^* from *P_PEPCK_, Δpepck* expressing *GDH2^His^* from *P_GDH2_*, *Δgdh2* expressing *PEPCK^Myc^* from *P_PEPCK_* and *Δaat2* expressing *PEPCK^Myc^* from *P_PEPCK_*

Expression cassette comprising of genes encoding PEPCK along with 1 kb of its promoter was cloned into *pIB3* vector (cat # 25452, Addgene, USA) and expressed in *P. pastoris GS115, Δrtg1*, *Δgdh2* and *Δaat2* as Myc-tagged proteins. *GDH2* was cloned with 545 bp of its promoter in *pIB3* vector as a His-tagged protein and expressed in *P. pastoris GS115, Δrtg1* and *Δpepck*. *PEPCK* was amplified from *GS115* genomic DNA using primer pair 5’ GG<u>GGTACC</u> CACCCACCCTCAAGTGC 3’ and 5’ CCC<u>AAGCTT</u>CTACAGGTCTTCTTCAGAGATC AGTTTCTGTTCC AACTGAGGGCCGGCCTG 3’ (KpnI and HindIII sites in the primers are underlined, respectively). GDH2 was amplified from *GS115* genomic DNA using primer pair 5’ CCG<u>GAATTC</u>CTCTCATGTTCGGATAATTCCAGCGGCTTTC 3’ and 5’ CCG<u>CTCGAG</u> CTAATGATGATGATGATGATGCAATCC CCGAGACTTGTAC 3’ (EcoRI and XhoI sites are underlined in the primers, respectively). PCR products were cloned into *pIB3* vector and transformed into *E. coli DH5α* competent cells. Recombinant plasmids containing *P_PEPCK_PEPCK-Myc* and *P_GDH2_GDH2-His* were linearized using BsrGI and StuI, respectively and transformed into respective *P. pastoris* strains by electroporation. Recombinant clones were selected by plating on YNBD His^−^ plates and clones expressing Myc-tagged PEPCK and His-tagged GDH2 were confirmed by western blotting using anti-Myc and anti-His antibodies.

### Generation of *P. pastoris GS115* and *Δrtg1* strains expressing GFP from *P_GDH2_* and *P_PEPCK_*

Gene encoding GFP was cloned under 1 kb promoter of *GDH2* and *PEPCK* into pIB3 vector (Addgene plasmid #25452) and expressed in *P. pastoris GS115* and *P. pastoris Δrtg1* (*Δrtg1*). For the generation of *P_GDH2_-GFP* construct, 1 kb *GDH2* promoter was amplified from *GS115* genomic DNA using primer pair 5’ CCGCTCGAGGGACAACCAAAGCATCC 3’ and 5’ CTC CTTTACTAGTCAGATCTACCATAGTGGGTTGGGAGTTTAGTGG 3’. The coding region of GFP was amplified from the vector, *pREP41GFP* (Craven et al., 1998) using primer pair 5’ CCACTAAACTCCC AACCCACTATGGTAGATCTGACTAGTAAAGGAG 3’ and 5’ CCCAAGCTTCTAGTGG TGGTGGCTAGCTTTG 3’. The amplified *GDH2* and *GFP* products were then purified and used as templates in the final PCR using primer pair 5’ CCG<u>CTCGAG</u>GGACAACCAAAGCATCC 3’ and 5’ CCC<u>AAGCTT</u>CTAGTGGTGGTGGCTAGCTTTG 3’ (XhoI and HindIII sites are underlined in the primers, respectively). The overlapping product was digested and ligated in *pIB3* vector followed by transformation in *E. coli DH5α* competent cells. For the generation of *P_PEPCK_-GFP* construct, 1 kb *PEPCK* promoter was amplified from *GS115* genomic DNA using primer pair 5’ GGGGTACCCACCCACCCTCAAGTGC 3’ and 5’ CCTTCTCATAG ATTATTATCCACAATGGTAGATCTGACTAGTAAAGGAG 3’. *GFP* was amplified using primer pair 5’ CTCCTTTACTAGTCAGATCTACCATTGTGGATAATAATCTATGAGAA GG 3’ and 5’ CCCAAGCTTCTAGTGGTGGTGGCTAGCTTTG 3’. The amplified *PEPCK* and *GFP* products were then purified and used as templates in the final PCR using primer pair 5’ GG<u>GGTACC</u>CACCCACCCTCAAGTGC 3’ and 5’ CCC<u>AAGCTT</u>CTAGTGGTGGTGGC TAGCTTTG 3’ (KpnI and HindIII sites are underlined in the primer respectively). The overlapping product was digested and ligated in *pIB3* vector followed by transformation in *E. coli DH5α* competent cells. Both the recombinant plasmids were linearised with SalI and transformed by electroporation into *GS115* and *Δrtg1*. Recombinant clones were selected by plating on YNBD His^−^ plates and clones expressing GFP were confirmed by western blotting using anti-GFP antibody.

### Generation of *GS115* strain expressing GFP from *P_AOX1_*

For the construction of *P_AOX1_-GFP*, 1 kb of AOX1 promoter was amplified by PCR from *P. pastoris* genomic DNA using the primer pair 5’ CGGG<u>GTACCT</u>CATGTTGGTATTGTGAA ATAGACGCAGATC 3’ and 5’ CTCCTTTACTAGTCAGATCTACCATCGTTTCGATAAT TAGTTGTTTTTTGATC 3’. KpnI site is underlined. GFP ORF was amplified from *pREP41GFP* (Craven et al., 1998) using the primer pair 5’ GATCAAAAAACAACTAATTATTCGAAACGATGG TAGATCTGACTAGTAAAGGAG 3’ and 5’ CCG<u>CTCGAG</u>CTAGTGGTGGTGGCTAGCT TTG 3’. XhoI site is underlined. In the third and final PCR reaction, the PCR products from the first two reactions were used as templates and amplified using 5’ CGGGGTACCTCATGTTGGTATTGTGAAATAGACGCAGATC 3’ and 5’ CCGCTCGAGCTAGTGGTGGTGGCTAGC TTTG 3’ primers to generate *P_AOX_-GFP* expression cassette, which was digested and cloned into *pIB3* to generate *pIB3-P_AOX_-GFP*. The recombinant plasmid was linearized with SalI, transformed into *GS115* and plated on YNBD His^−^ agar plates. Colonies expressing GFP were identified by western blotting of cell lysates using anti-GFP antibodies.

### Generation of *P. pastoris GS115* and *Δrtg1* strains expressing *GDH2^His^* and *PEPCK^Myc^* from *P_GAPDH_*

For the generation of *P_GAPDH_-GDH2^His^* and *P_GAPDH_-PEPCK^Myc^* expression cassettes, genes encoding *GDH2* and *PEPCK* were amplified from *GS115* genomic DNA and cloned into *pGHYB* vector (Addgene plasmid #87447) (Yang et al., 2014) under *GAPDH* promoter. For *GDH2* amplification 5’ CCG<u>CTCGAG</u>ATGGTCGACAGACTCCAAGTGTC 3’ and 5’ ATAAGAAT<u>GCGGCCGC</u> CAATCCCCGAGACTTGTACTC 3’ primer pair was used (XhoI and NotI sites are underlined respectively). For *PEPCK* amplification 5’ CCG<u>CTCGAG</u>ATGGCTCCTACTGCTATAGATTTAC 3’ and 5’ ATAAGAAT<u>GCGGCCGC</u>CTACAGGTCTTCTTCAGAGATCAGTTTCTGTTCCAACTG AGGGCCGGCCTG 3’ primer pair was used (XhoI and NotI sites are underlined, respectively). Both the recombinant plasmids were linearised with AvrII and transformed by electroporation into *GS115* and *Δrtg1*. Recombinant clones were selected by plating on hygromycin (200 μg/ml) containing YPD plates and clones expressing the epitope tagged proteins were confirmed by western blotting using anti-His and anti-Myc antibodies.

### Generation of *P. pastoris GS115* and *Δrtg1* strains expressing *GDH2^His^* and *PEPCK^Myc^* from *P_GAPDH_* containing *GDH2^DPR^* and *PEPCK^DPR^*

To generate *P_GAPDH-GDH2_^DPR^GDH2^His^* expression cassette, *GAPDH* promoter (Supplementary Fig. 1) from −1045 To −106, just upstream its TATAA box and *GDH2* from its putative TATAA box (−94) to stop codon were amplified from *GS115* genomic DNA. *GDH2* was amplifies with 6X-His tag sequence upstream its stop codon. *GAPDH* promoter was amplified using 5’ GGGGTACCGAAGTAAA ACTTTAACTTCAG 3’ and 5’ CTGGGCGATCTGGAGCTATT TATATTCGATTCTGGTG GTTTCCAATAATC 3’ primer pair. GDH2 was amplified using primer pair, 5’ GATTATTGGAAACCACCAGAATCGAATATAAATAGCTCCAGATCGC CCAG 3’ and 5’ CCGCTCGAGCTAATGATGATGATGATGATGCAATCCCCGAGACTT GTAC 3’. The PCR products were used as templates for final PCR using 5’ GG<u>GGTACC</u>GAA GTAAAACTTTAACTTCA G 3’ and 5’ CCG<u>CTCGAG</u>CTAATGATGATGATGATGATGC AATCCCCGAGACTTGTA C 3’ primer pair (KpnI and XhoI sites are underlined respectively) and cloned into *pIB3* vector (cat # 25452, Addgene, USA). Similar strategy was used for *P_GAPDH-PEPCK_^DPR^PEPCK^Myc^* construct generation. Primer pair 5’ GG<u>GGTACC</u>GAAGTAAAA CTTTAACTTCAG 3’ and 5’ GGCGGCCAGCCTGCCCCTATTTATATTCGATTCTGGTGGTTTCCAATAATC 3’ was used for *GAPDH* promoter (from −1045 to −106, just upstream its TATA box) amplification and primer pair 5’ GATTATTGGAAACCACCAGAATCGAATATAAATAGGGGCAGGCTGGCCGCC 3’ and 5’ CCC<u>AAGCTT</u>CTACAGGTCTTCTTCA GAGATCAGTTTCTGTTCCAACTGAGGGCCGGCCTG 3’ was used for *PEPCK* (from −66 bp to stop codon) amplification. Final PCR was done to generate the desired construct using primer pair 5’ GG<u>GGTACC</u>GAAGTAAAACTTTAACTTCAG 3’ and 5’ CCC<u>AAGCTT</u>CTA CAGGTCTTCTTCAGAGATCAGTTTCTGTTCCAACTGAGGGCCGGCCTG 3’ (KpnI and HindIII sites are underlined respectively). Both the recombinant plasmids were linearised with StuI and transformed by electroporation into *GS115* and *Δrtg1*. Recombinant clones were selected by plating on YNBD His^−^ plates and clones expressing the epitope tagged proteins were confirmed by western blotting using anti-His and anti-Myc antibodies.

### GST Pull down assay

*E. coli* cells (*BL21DE3*) expressing GST tagged anti-GFP nanobody (Addgene plasmid #61838) (Katoh et al., 2015) were suspended in 1X PBS (137 mM NaCl, 2.7 mM KCl, 100 mM Na_2_HPO_4_ and 2 mM KH_2_PO_4_, pH-7.4) containing 1 mM PMSF, 1 mM EDTA, 1 mM DTT, 10μg/ml lysozyme and 1% Triton X-100 and kept on ice for 20 min. Cells were then sonicated and cell lysate containing GST tagged anti-GFP nanobody was incubated with glutathione resin (G-Biosciences, U.S.A.) at 4°C for 1 h. Nanobody bound glutathione resins were harvested by brief centrifugation followed by washes with 1X PBS containing 1 mM PMSF. GST tagged anti-GFP nanobody bound glutathione resins were incubated with yeast (*GS115*) whole cell protein lysate containing GFP expressing from either *P_AOX1_, P_GDH2_* or *P_PEPCK_* at 4°C for 2-3 h. Post interaction resins were centrifuged briefly and washed atleast thrice with cold 1X PBS. Bound proteins were resolved by SDS-PAGE and visualized by Coomassie Brilliant Blue R staining.

### Statistical analysis

The relevant statistical test and replicate type for each figure are found in the corresponding figure legends.

## Data availability

The authors declare that all data supporting the findings of the present study are available in the article and its supplementary figures and tables, or from the corresponding author upon request.

## Acknowledgements

We thank Kamisetty Krishna Rao for providing *P. pastoris GS115* expressing *P_AOXI_-GFP*. This work was supported by the research grant EMR/2015/000567 and J. C. Bose Fellowship grant SB/S2/JCB-025/2015 awarded by the Science and Engineering Research Board, New Delhi, India and the research grant BT/PR30986/BRB/10/1751/2018 awarded by the Department of Biotechnology, New Delhi, India (to P.N.R). Funding from the Department of Science and Technology Fund for Improvement of S&T Infrastructure in Higher Educational Institutions (DST-FIST), the University Grants Commission and the Department of Biotechnology (DBT)-Indian Institute of Science partnership program is acknowledged. The Senior Research Fellowship to T.D. from the Department of Biotechnology, New Delhi is gratefully acknowledged.

## Author contributions

T. Dey designed and performed experiments and analyzed the data. P. N. R. conceived the project, designed experiments and analyzed the data. T.D. and P.N.R. wrote the manuscript.

## References

Adachi, A., M. Koizumi, and Y. Ohsumi. 2017. Autophagy induction under carbon starvation conditions is negatively regulated by carbon catabolite repression. J. Biol. Chem. 292:19905–19918. http://doi.org/10.1074/jbc.M117.817510.

Butow, R.A., and N.G. Avadhani. 2004. Mitochondrial signaling: The retrograde response. Mol. Cell. 14:1–15. http://doi.org/10.1016/S1097-2765(04)00179-0.

Cereghino, J.L., and J.M. Cregg. 2000. Heterologous protein expression in the methylotrophic yeast *Pichia pastoris*. FEMS Microbiol. Rev. 24:45–66. http://doi.org/10.1016/S0168-6445(99)00029-7.

Craven, R.A., D.J.F. Griffiths, K.S. Sheldrick, R.E. Randall, I.M. Hagan, and A.M. Carr. 1998. Vectors for the expression of tagged proteins in *Schizosaccharomyces pombe*. Gene. 221:59–68. http://doi.org/10.1016/S0378-1119(98)00434-X.

Dey, T., K.K. Rao, J. Khatun, and P.N. Rangarajan. 2018. The nuclear transcription factor Rtg1p functions as a cytosolic, post-transcriptional regulator in the methylotrophic yeast *Pichia pastoris*. J. Biol. Chem. 293:16647–16660. http://doi.org/10.1074/jbc.RA118.004486.

Freese, S., T. Vogts, F. Speer, B. Schäfer, V. Passoth, and U. Klinner. 2011. C-and N- catabolic utilization of tricarboxylic acid cycle-related amino acids by *Scheffersomyces stipitis* and other yeasts. Yeast. 28:375–390. http://doi.org/10.1002/yea.1845.

Garst, A.D., A.L. Edwards, and R.T. Batey. 2011. Riboswitches: structures and mechanisms. Cold Spring Harb. Perspect. Biol. http://doi.org/10.1101/cshperspect.a003533.

Hartner, F.S., and A. Glieder. 2006. Regulation of methanol utilisation pathway genes in yeasts. Microb. Cell Fact. 5:1–21. http://doi.org/10.1186/1475-2859-5-39. https://www.amazon.in/Ajinomoto-Monosodium-Glutamate-Umami-Seasoning/dp/B00IH28XDE. https://www.fda.gov/food/food-additives-petitions/questions-and-answers-monosodium-glutamate-msg. https://www.yeastgenome.org/locus/S000002374#protein.

Iwama, R., and Y. Ohsumil. 2019. Analysis of autophagy activated during changes in carbon source availability in yeast cells. J. Biol. Chem. 294:5590–5603. http://doi.org/10.1074/jbc.RA118.005698.

Jazwinski, S.M. 2014. The retrograde response: a conserved compensatory reaction to damage from within and from without. Prog. Mol. Biol. Transl. Sci. 127:133–154. https://doi.org/10.1016/B978-0-12-394625-6.00005-2.

Katoh, Y., S. Nozaki, D. Hartanto, R. Miyano, and K. Nakayama. 2015. Architectures of multisubunit complexes revealed by a visible immunoprecipitation assay using fluorescent fusion proteins. J. Cell Sci. 128:2351–2362. http://doi.org/10.1242/jcs.168740.

Magasanik, B. 1993. Regulation of nitrogen utilization. COLD SPRING Harb. Monogr. Ser. 21:283. http://dx.doi.org/10.1101/0.283-317.

Mara, P., G.S. Fragiadakis, F. Gkountromichos, and D. Alexandraki. 2018. The pleiotropic effects of the glutamate dehydrogenase (GDH) pathway in *Saccharomyces cerevisiae*. Microb. Cell Fact. 17:170. http://doi.org/10.1186/s12934-018-1018-4.

McMillan, J., Z. Lu, J.S. Rodriguez, T.-H. Ahn, and Z. Lin. 2019. YeasTSS: an integrative web database of yeast transcription start sites. Database. 2019.

Miyashita, Y., and A.G. Good. 2008. NAD(H)-dependent glutamate dehydrogenase is essential for the survival of *Arabidopsis thaliana* during dark-induced carbon starvation. J. Exp. Bot. 59:667–680. http://doi.org/10.1093/jxb/erm340.

Prielhofer, R., M. Maurer, J. Klein, J. Wenger, C. Kiziak, B. Gasser, and D. Mattanovich. 2013. Induction without methanol: novel regulated promoters enable high-level expression in *Pichia pastoris*. Microb. Cell Fact. 12:1–10. http://doi.org/10.1186/1475-2859-12-.

Sahu, U., and P.N. Rangarajan. 2016. Methanol expression regulator 1 (Mxr1p) is essential for the utilization of amino acids as the sole source of carbon by the methylotrophic yeast, *Pichia pastoris*. J. Biol. Chem. 291:20588–20601. http://doi.org/10.1074/jbc.M116.740191.

Schmitt, M.E., T.A. Brown, and B.L. Trumpower. 1990. A rapid and simple method for preparation of RNA from *Saccharomyces cerevisiae*. Nucleic Acids Res. 18:3091–3092. http://doi.org/10.1093/nar/18.10.3091.

Shen, W., Y. Xue, Y. Liu, C. Kong, X. Wang, M. Huang, M. Cai, X. Zhou, Y. Zhang, and M. Zhou. 2016. A novel methanol-free *Pichia pastoris* system for recombinant protein expression. Microb. Cell Fact. 15:1–11. http://doi.org/10.1186/s12934-016-0578-4.

Silao, F.G.S., K. Ryman, T. Jiang, M. Ward, N. Hansmann, C. Molenaar, N.-N. Liu, C. Chen, and P.O. Ljungdahl. 2020. Glutamate dehydrogenase (Gdh2)-dependent alkalization is dispensable for escape from macrophages and virulence of *Candida albicans*. PLoS Pathog. 16:e1008328. https://doi.org/10.1371/journal.ppat.1008328.

Miller, S.M., and B. Magasanik. 1990. Role of NAD-linked glutamate dehydrogenase in nitrogen metabolism in *Saccharomyces cerevisiae*. J. Bacteriol. 172:4927–4935. http://doi.org/10.1128/jb.172.9.4927-4935.1990

Sutter, B.M., X. Wu, S. Laxman, and B.P. Tu. 2013. Methionine Inhibits Autophagy and Promotes Growth by Inducing the SAM-Responsive Methylation of PP2A. Cell. 154:403–415. https://doi.org/10.1016/j.cell.2013.06.041.

Vogl, T., L. Sturmberger, P.C. Fauland, P. Hyden, J.E. Fischer, C. Schmid, G.G. Thallinger, M. Geier, and A. Glieder. 2018. Methanol independent induction in *Pichia pastoris* by simple derepressed overexpression of single transcription factors. Biotechnol. Bioeng. 115:1037–1050. http://doi.org/10.1002/bit.26529.

Wang, J., X. Wang, L. Shi, F. Qi, P. Zhang, Y. Zhang, X. Zhou, Z. Song, and M. Cai. 2017. Methanol-Independent Protein Expression by AOX1 Promoter with trans-Acting Elements Engineering and Glucose-Glycerol-Shift Induction in *Pichia pastoris*. Sci. Rep. 7:1–12. http://doi.org/10.1038/srep41850.

Weber, C.A., K. Sekar, J.H. Tang, P. Warmer, U. Sauer, and K. Weis. 2020. β-Oxidation and autophagy are critical energy providers during acute glucose depletion in *Saccharomyces cerevisiae*. Proc. Natl. Acad. Sci. 117:12239 LP–12248. http://doi.org/10.1073/pnas.1913370117.

Yang, J., L. Nie, B. Chen, Y. Liu, Y. Kong, H. Wang, and L. Diao. 2014. Hygromycin-resistance vectors for gene expression in *Pichia pastoris*. Yeast. 31:115–125. http://doi.org/10.1002/yea.3001.

