## Supplementary file for "Post-transcriptional regulation of glutamate metabolism of *Pichia pastoris* and development of a glutamate-inducible yeast expression system"

**Supplementary data**

**Supplementary Fig.1**

**a**. *P. pastoris GDH2* promoter (-1000) Locus tag/Gene: PAS_chr2-1_0311

cttttggacaaccaaagcatccgcctgaagggacaacttgcattcaacggcttcagttggaaacgtcagagctgacctatagtttgctaga
accgttttctctgtttacgtttacgtctcctcaaatttgcgctcggtatgtccttcctaattagcgggaaaagctgttcttagttaatacggagaa
agtttcggggttaccgttccgggaagaggaggggtcatctctctcatctcatccaaccattaagtttcttccaaaacttcaggataatcagttt
aaccaccgacaggagtcagatttgagattgacagaaagtttttccgtccatttcctcatcttgtcgccgttatcagtcaatctctatggttatct
ggaatttctttttcttttaattcatcttctttttatcccgcgcctttggcgttctagctcatctcatgaaaacaaaaccctctcatgttcggataattc
cagcggctttcactttcagatgacacatagattggactcaaccatggctatctggggtatacggacgttggcaagggcgttaatttttcagg
acaaacggaaatgccatggctccagggaaaggcattcctattgcaaacctagaccgtcgaacctctcctatcgcctaccagtcacccagc
tatccctaggcaactcatctccttcaagcggattgcaacctgctaagccaaattagatctggccacagaaatgccgcaatatttcttggctct
cccctccctctggtcttctctcattatcttgcatccttctggcattgtggttttcagaggtttttctaaaaaaataatcacgaccgatggggtctcg
atgaggcatctcttaagtcatttttttcccccacaaacagaaaaagcaaaaaactccgtataaatagctccagatcgcccagacgaatctcg
cccttgactcaattagtagaaagaaactaaaactaccccccactaaactcccaacccactATG
-1

**b.** *P. pastoris PEPCK* promoter (-1000) Locus tag/Gene: PAS_FragB_0061

ctacatcggaaccaattttgaattcgccgatacgaaaccggacatcaagttccatcatcatgcaagatgatcatcaggaagagccgtggtc
agtcaacactgcaagtgtcaaagagacatcaattattgacattagtaccccggatgaggcttacaatgctcgcgggaaagacgatgcgat
acgggttgattcggagggcccttcaaaaagaacaactattctcagccgtctgatggagacacaagattcggacaatgacggaacggagt
ctgacgtaggggagagttctagtctgataattcttagttgaccagccttgcatcgggttgaacaaacttttgtgggttgggccactttgcgcc
aacaacacaaggttttcacaccacccggttgtcattaccgcccggaaaactcaatgattgacggagcgtaggttcacggaagtcaagtta
gagtagtcggagagtttcgggtgataagaatttcccgggtgctttggagtgccaaagtgactgaatcaggaactaaaaacccgggctaac
ggctgaaggcctgtgttcagtgcactatattgtattctagtctggggtaggttgaagtttggtggattcacgactccggagtgatggaggga
caataccgaagatgaggtcttgcgagtggaaagataggttaagaataattaataacattgtatgaaacaagatggggagacaattttagct
atcgatccgttatcattggtcgaatgatgacttggaaccttatttgcactttttttgtcaggtgttcgccgaaacttaaagatagttttattgatgtg
tgccaaaaatgtggggacaaaaggtcctccacccgccatcccccggtctcatacaaaaaaaaagaaaaatacctctactagtccggggt
caaggcaagtcatgataggcatataaataggggcaggctggccgccattacgaattagattaaccttctcatagattattatccacaATG
-1

**c**. *P. pastoris GAPDH* promoter (-1048) Locus tag/Gene: PAS_chr2-1_0437

tttgaagtaaaactttaacttcagctccttacatttgcactaagatctctgctactctggtcccaagtgaaccaccttttggaccctattgaccg
gaccttaacttgccaaacctaaacgcttaatgcctcagacgttttaatgcctctcaacacctccaaggttgctttcttgagcatgcctactagg
aactttaacgaactgtggggttgcagacagtttcaggcgtgtcccgaccaatatggcctactagactctctgaaaaatcacagttttccagta
gttccgatcaaattaccatcgaaatggtcccataaacggacatttgacatccgttcctgaattatagtcttccaccgtggatcatggtgttcctt
tttttcccaaagaatatcagcatcccttaactacgttaggtcagtgatgacaatggaccaaattgttgcaaggtttttctttttctttcatcggcac
atttcagcctcacatgcgactattatcgatcaatgaaatccatcaagattgaaatcttaaaattgcccctttcacttgacaggatcctttttttgta
gaaatgtcttggtgtcctcgtccaatcaggtagccatctctgaaatatctggctccgttgcaactccgaacgacctgctggcaacgtaaaatt
ctccggggtaaaacttaaatgtggagtaatggaaccagaaacgtctcttcccttctctctccttccaccgcccgttaccgtccctaggaaatt
ttactctgctggagagcttcttctacggcccccttgcagcaatgctcttcccagcattacgttgcgggtaaaacggaggtcgtgtacccgac
ctagcagcccagggatggaaaagtcccggccgtcgctggcaataatagcgggcggacgcatgtcatgagattattggaaaccaccaga
atcgaatataaaaggcgaacacctttcccaattttggtttctcctgacccaaagactttaaatttaatttatttgtccctatttcaatcaattgaaca
actatcaaaacacaATG
-1

**Fig 1. Sequence and locus tag/gene IDs of *P. pastoris* promoters used in this study. a)** Nucleotide sequence of -1000bp of *GDH2* promoter. **b)** Nucleotide sequence of -1000bp of *PEPCK* promoter. **c)** Nucleotide sequence of -1048bp of *GAPDH* promoter. Putative TATA box is underlined. Initiation codon (ATG) is shown in upper case.

**Supplementary Table 1**

*P. pastoris* strains used in this study

| Yeast strain (*P. pastoris*) | Description | Source |
|---|---|---|
| *GS115* | *his4* | Ref. (4) |
| *Δrtg1* | *GS115, rtg1Δ ::Zeo$^r$* | Ref. (2) |
| *Δgdh2* | *GS115, gdh2Δ ::Zeo$^r$* | Ref. (1) |
| *Δpepck* | *GS115, pepckΔ ::Zeo$^r$* | Ref. (2) |
| *Δaat2* | *GS115, aat2Δ ::Zeo$^r$* | Ref. (1) |
| *GS115-$P_{GDH2}$-GDH2$^{His}$* | *GS115, his$^{4+}$:: (pIB3-$P_{GDH2}$-GDH2-His)* | Ref. (1) |
| *GS115-$P_{PEPCK}$-PEPCK$^{Myc}$* | *GS115, his$^{4+}$:: (pIB3- $P_{PEPCK}$-PEPCK-Myc)* | This study |
| *GS115-$P_{AOX1}$-GFP* | *GS115, his4$^+$:: (pIB3-$P_{AOX1}$GFP)* | This study |
| *GS115-$P_{GDH2}$-GFP* | *GS115, his4$^+$:: (pIB3-$P_{GDH2}$GFP)* | This study |
| *Δrtg1-$P_{GDH2}$-GFP* | *Δrtg1, his4$^+$:: (pIB3-$P_{GDH2}$GFP)* | This study |
| *GS115-$P_{PEPCK}$-GFP* | *GS115, his4$^+$:: (pIB3-$P_{PEPCK}$GFP)* | This study |
| *Δrtg1-$P_{PEPCK}$-GFP* | *Δrtg1, his4$^+$:: (pIB3-$P_{PEPCK}$GFP)* | This study |
| *GS115-$P_{GAPDH}$-GDH2$^{His}$* | *GS115, Hyg$^r$:: (pGHYB-$P_{GAPDH}$-GDH2-His)* | This study |
| *Δrtg1-$P_{GAPDH}$-GDH2$^{His}$* | *Δrtg1, Hyg$^r$ :: (pGHYB-$P_{GAPDH}$-GDH2-His)* | This study |
| *GS115-$P_{GAPDH}$-PEPCK$^{Myc}$* | *GS115, Hyg$^r$:: (pGHYB-$P_{GAPDH}$-PEPCK-Myc)* | This study |
| *Δrtg1-$P_{GAPDH}$-PEPCK$^{Myc}$* | *Δrtg1, Hyg$^r$ :: (pGHYB-$P_{GAPDH}$-PEPCK-Myc)* | This study |
| *GS115-$P_{GAPDH\text{-}GDH2}{}^{DPR}$-GDH2$^{His}$* | *GS115, his4$^+$ :: (pIB3-$P_{GAPDH\text{-}GDH2}{}^{DPR}$-GDH2-His)* | This study |
| *Δrtg1-$P_{GAPDH\text{-}GDH2}{}^{DPR}$-GDH2$^{His}$* | *GS115, his4$^+$ :: (pIB3-$P_{GAPDH\text{-}GDH2}{}^{DPR}$-GDH2-His)* | This study |
| *GS115-$P_{GAPDH\text{-}PEPCK}{}^{DPR}$-PEPCK$^{Myc}$* | *GS115, his4$^+$ ::(pIB3-$P_{GAPDH\text{-}PEPCK}{}^{DPR}$-PEPCK-Myc)* | This study |
| *Δrtg1-$P_{GAPDH\text{-}PEPCK}{}^{DPR}$-GDH2$^{His}$* | *GS115, his4$^+$ ::(pIB3-$P_{GAPDH\text{-}PEPCK}{}^{DPR}$-PEPCK-Myc)* | This study |
| *Δpepck-$P_{GDH2}$-GDH2$^{His}$* | *Δpepck, his$^{4+}$:: (pIB3-$P_{GDH2}$-GDH2-His)* | This study |
| *Δgdh2-$P_{PEPCK}$-PEPCK$^{Myc}$* | *Δgdh2, his$^{4+}$:: (pIB3- $P_{PEPCK}$-PEPCK-Myc)* | This study |
| *Δaat2-$P_{PEPCK}$-PEPCK$^{Myc}$* | *Δaat2, his$^{4+}$:: (pIB3- $P_{PEPCK}$-PEPCK-Myc)* | This study |
